# Structural Basis for Dimeric Genome Selection by HIV-1

**DOI:** 10.64898/2026.06.17.732839

**Authors:** Nele Merret Hollmann, Siarhei Kharytonchyk, Angela Buchannan, Caela Phillips, Amir Ledbetter, Fairine Ahmed, Anna Adelstein, Kaylee Barry, Samuel A. Geleta, Cleo Burnett, Alice Telesnitsky, Barbie K. Ganser-Pornillos, Owen Pornillos, Michael F. Summers

## Abstract

HIV-1 selects and packages a dimeric RNA genome into assembling virions, a requirement for strand transfer-mediated recombination during reverse transcription. Conserved RNA elements within the 5’-leader that promote dimerization and packaging have been identified (dimer initiation signal, DIS; core encapsidation signal, Ψ^CES^; respectively), but the molecular mechanism that couples these processes remains unknown. Using an in vitro virus assembly assay, we show that dimeric HIV-1 5’-Leader RNAs promote assembly of viral Gag proteins into protein lattices resembling those of immature virions, whereas monomeric 5’-Leader constructs with similar affinities for Gag do not. Assembly requires the cellular factor, inositol hexakisphosphate, mimicking biology. Cryogenic electron microscopy (cryoEM) and tomography (cryoTM) studies of nascent assemblies are consistent with a structural model in which the dimeric DIS_2_ helix bridges two elongated 6-helix bundles formed by the SP1 domains of adjacent Gag hexamers. The structure positions clusters of RNA-binding zinc knuckles from Gag’s nucleocapsid domain in close proximity to clusters of conserved guanosines shown here to be essential for genome packaging. Our findings suggest that DIS_2_ functions as a molecular ruler, organizing Gag hexamers into structures optimized for Gag recruitment and Gag lattice expansion.

## INTRODUCTION

Like most retroviruses, human immunodeficiency virus type-1 (HIV-1) packages two copies of its RNA genome (gRNA) into assembling virus particles 1-5. Both RNA strands are required for replication, enabling strand-transfer mediated recombination during reverse transcription to overcome breaks in the fragile genome and to promote genetic diversity 6. Genomes are recognized by the nucleocapsid domains (NC) of the viral Gag protein and anchored to the plasma membrane by a small number of Gag molecules (several dozen or fewer) 7,8. The resulting ribonucleoprotein complex then serves as a nucleant for virus assembly, recruiting several thousand additional Gag proteins and cellular factors required for virus budding and release 9. Genomes are packaged as non-covalently linked dimers, and overlapping RNA elements that promote dimerization and packaging (dimer initiation signal, DIS; core encapsidation signal, Ψ^CES^, respectively) reside within the ∼350 nucleotide 5’-leader of the viral transcript (Fig. 1a) 2,10-13, The dimeric form of the HIV-1 5’-leader exposes approximately 32 NC binding sites (20 with Kd values of ∼45 to 330 nM) whereas the monomeric form possesses only ∼7 high affinity NC binding sites 14. These and other studies are consistent with proposals that RNA dimerization and packaging are mechanistically coupled 1,4,5,12,13,15-22.

**Figure 1.**
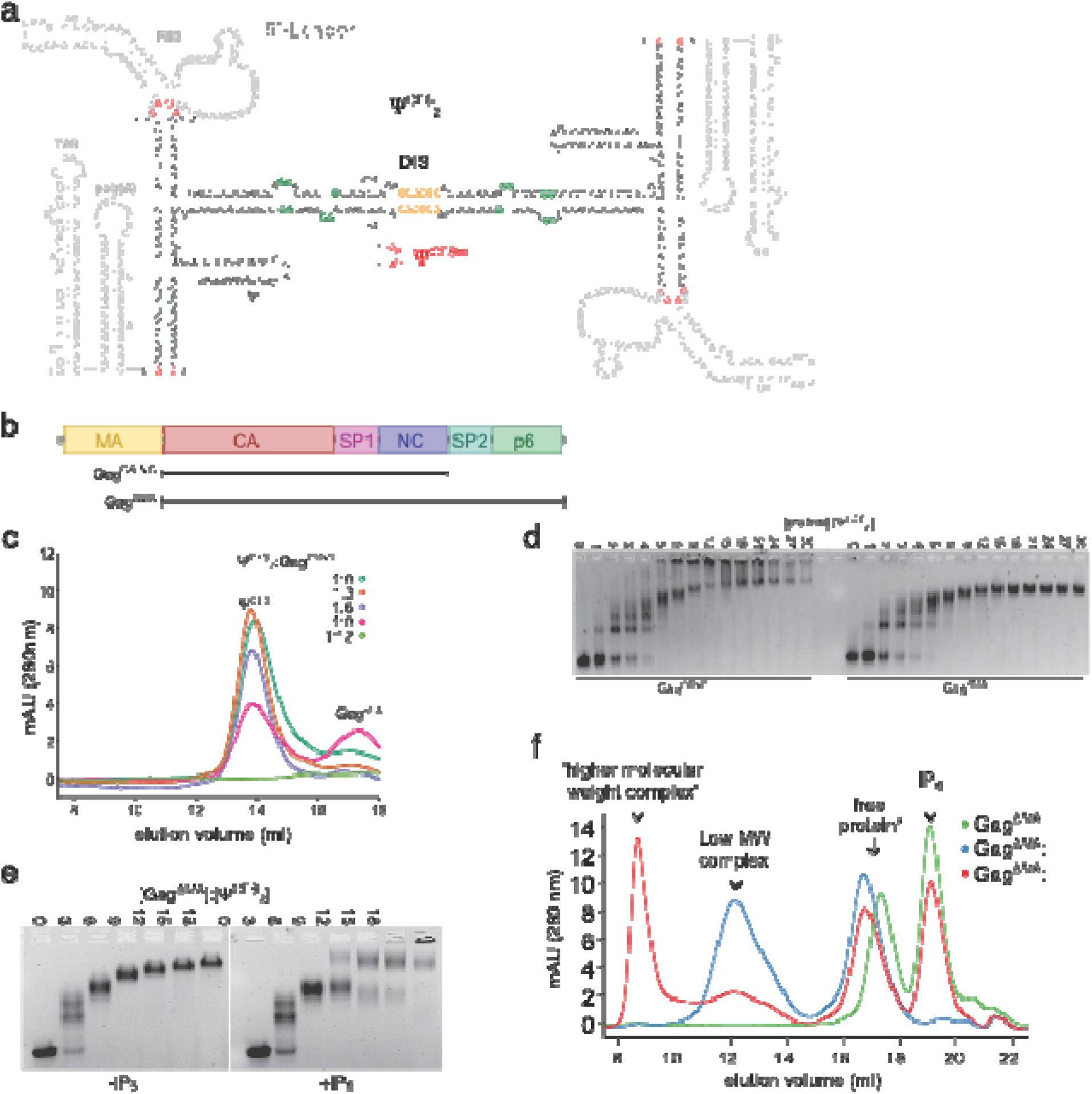
A soluble HIV-1 Gag:Ψ^CES^_2_ complex resembling IP_6_ dependent lattice formation. **a:** Secondary structure of the dimeric 5’ leader RNA. Introduced mutations are highlighted in red. Nucleotides belonging to the Ψ^CES^ are shown in black. The palindromic sequence nucleating dimerization is shown in yellow. Exposed guanosines important for nucleocapsid binding are highlighted green. **b:** Schematic representation of HIV-1 Gag protein constructs (Gag^CA-SP2-NC^, Gag^ΔMA^). **c:** Analytical SEC data of complex formation at different Gag^CA-SP2-NC^:Ψ^CES^_2_ ratios, indicating the formation of an insoluble complex (teal: 1:0; orange 1:3; violet: 1:6; pink: 1:9; green: 1:12). **d:** A gel shift assay compares the influence of SP1-p6 addition on ribonucleoprotein complex formation. The absence of complex trapped in the gel pocket for the Gag^ΔMA^ constructs indicates an increased solubility of the complex. Increasing Gag: Ψ^CES^_2_ ratios from 1:0 up to 1:30 are shown. **e:** Gel shift assay showing the cooperative effect of IP_6_ addition on Gag^ΔMA^: Ψ^CES^_2_ complex formation. **f:** Analytical SEC demonstrating the formation of two distinct complexes. A low molecular weight complex between Gag^ΔMA^ and Ψ^CES^_2_ is observed (blue). Upon addition of IP_6_ (1 mM), a heavier complex forms (red). No complex is detected in the absence of RNA (green). The protein:RNA ratio is 1:12. Complex samples were crosslinked with 0.01% Glutaraldehyde leading to a shift for the free protein due to the dimer interface in the CA domain.

Genome dimerization and NC binding are modulated at the level of transcription by heterogeneous start site (HTSS) usage 23,24. 5’-capped RNAs transcribed with a single 5’-guanosine preferentially form dimers in vitro and are selected for packaging as gRNAs whereas those transcribed with two or three 5□-guanosines adopt a monomeric leader structure in vitro and are retained in cells as mRNAs 24,25. The additional guanosines induce structural remodeling that exposes the 5’-cap and RNA elements important for splicing and translation, and simultaneously sequesters NC binding sites 14,25. These findings support a mechanism in which dimerization promotes packaging by both promoting NC binding and preventing dominant negative cap-dependent capture by the cellular RNA processing and translation machinery 26. An unexplained caveat is that mutations that promote co-transcriptional shedding of the 5’-cap and also block Ψ^CES^ RNA dimerization without affecting the secondary structure or NC binding properties of the RNA (Ψ^CESm^ ) 27 are poorly packaged 26. This indicates that cap sequestration and NC binding site exposure alone do not account for wild-type RNA packaging efficiency 26.

Recent in vitro virus assembly studies suggest that Gag:Gag interactions may also play a role in RNA packaging 28-33. The CA-SP1 junction of Gag promotes assembly of the immature lattice by adopting an intermolecular 6-helix bundle (6HB) stabilized by a centrally coordinated inositol-hexakisphosphate (IP6) molecule 34-40, and the C-terminal domain of CA (CACTD) forms a dimer interface that is essential for extending the Gag lattice 34,35,41,42. The presence of either nucleic acids or IP6 can promote Gag lattice formation with relatively slow kinetics 28,30,38,40, whereas inclusion of both RNA and IP6 can accelerate assembly 33. In all these studies, Gag assembles into spherical virus-like particles (VLP) comprising more than a thousand Gag proteins. Although detailed structural information has been obtained for CA lattices in immature virions and CA-containing assemblies, structural information for the RNA and its interactions with Gag in viral assemblies is lacking 34,40,43-45. Here we report an in vitro system for preparation of smaller, soluble Gag:RNA species amenable to characterization by cryoEM. Assembly is dependent on IP_6_ and was observed for both the dimeric Ψ^CES^ (but not a monomeric analog) and for a shorter dimeric DIS construct (DIS_2_; residues 237 – 281, Fig. 1a). Although cryoEM densities attributed to the RNA were not sufficient for high resolution analysis, the data point to a key role for the dimeric DIS_2_ element in scaffolding Gag assembly.

## RESULTS AND DISCUSSION

### Gag constructs and conditions that promote virus-like lattice assembly

A series of Gag mutants was prepared including chimeric proteins with domains engineered to stabilize dimeric, trimeric or hexameric Gag interfaces, and their solubility and assembly properties upon titration with Ψ^CES^_2_ and /or IP6 were assessed. A Gag construct lacking the MA, SP2 and p6 domain of Gag (GagCA-SP1-NC) (Fig. 1b), which is similar to a previously employed construct that lacked most residues of MA and was truncated at NC 28,30,33, formed insoluble precipitates under the conditions employed (20 mM MOPS buffer, 140 mM KCl, 10 mM NaCl, 5 mM TCEP, pH 7.5, [RNA] > 0.5 μM) with no detectable soluble RNA when Ψ^CES^_2_:Gag complexes were formed at ratios exceeding 1:6 (Fig. 1c). Similar results were obtained for a large number of Gag-derived protein constructs and conditions tested (Fig. S1). However, one construct lacking only the MA domain (but retaining the C-terminal SP2p6 regions; Gag^ΔMA^) remained soluble upon addition of Ψ^CES^_2_ (Fig. 1d). In the absence of IP6, binding occurred without apparent cooperativity, reaching saturation at a Gag^ΔMA^:RNA ratio of ∼6:1. Negative-stain EM revealed that these assemblies adopt heterogeneous, dumbbell-shaped structures with dimensions similar to that of an NMR-derived Ψ^CES^_2_ model (Fig. S2a). Conformational heterogeneity has thus far precluded further structural analyses of these low-MW complexes. However, based on the negative stain imaging and lack of binding cooperativity, it appears that Gag^ΔMA^ does not assemble into a regular lattice when binding to Ψ^CES^_2_.

Since IP6 is known to bind the 6HB of the immature Gag lattice, we evaluated its influence on Ψ^CES^_2_ mediated Gag^ΔMA^ assembly. In the presence of IP6, Gag^ΔMA^:Ψ^CES^_2_ assembly was markedly altered, triggering cooperative Gag^ΔMA^ binding and producing higher molecular weight yet soluble assemblies (Fig. 1e, f). Negative-stain EM imaging revealed assemblies with hexameric features resembling those of well-characterized immature CA lattices (Fig. S2b) 34,44. Soluble assemblies of various sizes were detected, with apparent surface areas ranging from 400 to 4000 nm2 (Fig. S2b – stained images). Previous studies showed that mutations in the DIS that block Ψ^CES^ dimerization without affecting its structure or NC binding properties inhibit packaging into virions 27.

To assess the role of RNA dimerization in Gag assembly, studies were conducted with a Ψ^CES^ mutant in which the dimer promoting XXGCGCGCXX residues (G = guanosine, C = cytosine, X = G, C, adenosine (A) or uridine (U) nucleotides) were substituted by a GAGA tetraloop (Ψ^CESm^) (Fig. 1a). Under conditions in which Ψ^CES^_2_ promotes assembly, the monomeric RNA formed only lower molecular weight complexes (Fig. S3a). Titrations with other RNAs, including tRNA, the Ψ-stem loop that binds NC with high affinity 46, and NTPs, yielded no higher molecular weight assemblies (Fig. 2a, Fig. S3b). These results indicate that the presence of high affinity NC binding sites alone is insufficient to promote Gag^ΔMA^ assembly under the assembly conditions employed, even when IP6 is included in the assembly assay.

**Figure 2.**
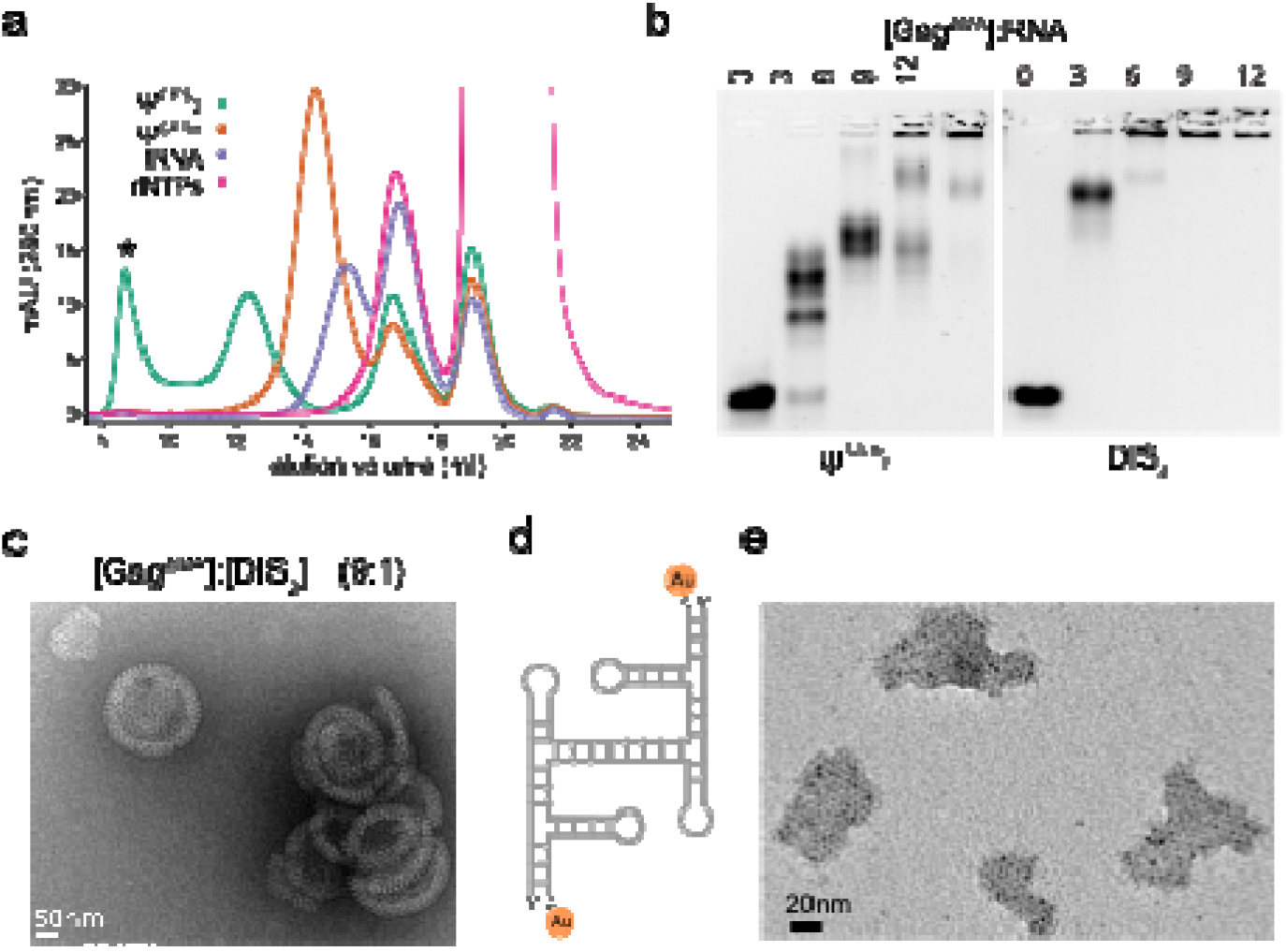
RNA characterization within the Gag^ΔMA^: Ψ^CES^_2_ complex revealed DIS_2_ necessity to assemble capsid lattice. **a:** Assembly formation is dependent on Ψ^CES^_2_ (teal). Neither a monomeric mutant Ψ^CESm^ (orange), tRNA (violet) nor the addition of NTPs (pink) results in the formation of higher-order assemblies. The protein:RNA ratio is 1:12. Only Ψ^CES^_2_ supports the formation of higher molecular weight assemblies (denoted * ) in the presence of IP_6_ (1 mM). **b:** A gel shift assay showing differences in complex formation between Ψ^CES^_2_ and DIS_2_. DIS_2_ is fully bound by Gag^ΔMA^ at a 3x molar excess, while free RNA remains at the same ratio for Ψ^CES^_2_. Insoluble aggregates form for DIS_2_ at 6x molar excess, whereas Ψ^CES^_2_ requires a 1:9 ratio. **c:** Negative-stain EM at a 1:9 ratio reveals that DIS_2_ and Gag^ΔMA^ form nearly fully-assembled immature virion-like capsules. **d:** Schematic representation of Ψ^CES^_2_ showing the positioning of a 3’ ligated gold nanoparticle. **e:** Negative stain EM of 3’ gold-labeled Ψ^CES^_2_ (left) demonstrates that RNA molecules are distributed throughout the assemblies (right). Assemblies were formed at a protein:RNA ratio of 1:9. Scale bar 20nm.

Since cooperative assembly requires the dimeric state of Ψ^CES^, we evaluated assembly in the presence of a shorter RNA construct comprising only the DIS_2_ duplex. This construct includes several non-base paired guanosine residues previously implicated to be important for RNA packaging 27. As observed for Ψ^CES^_2_, DIS_2_ exhibited IP6-dependent cooperative binding to Gag^ΔMA^. However, DIS_2_ induced assembly at lower protein:RNA ratio (1:3 vs. 1:6 for Ψ^CES^_2_), and larger assemblies appeared at lower protein concentrations (1:6 vs. 1:9) (Fig. 2b). Negative stain EM imaging confirmed that at a 1:3 RNA:Gag^ΔMA^ ratio, Ψ^CES^_2_ produced only sparse hexameric lattices whereas DIS_2_ yielded abundant assemblies (Fig. S4). DIS_2_ also induced formation of nearly completely formed virus-like particles (Fig. 2c) under conditions where Ψ^CES^_2_ produced much smaller, partial assemblies. These findings indicate that the DIS_2_ duplex alone is sufficient to promote IP6-dependent Gag lattice assembly in vitro, and does so with enhanced efficiency compared to that of the full-length, dimeric Ψ^CES^_2_ RNA.

### Cryo-EM reveals RNA-associated density bridging adjacent Gag hexamers

Although spectrophotometric 260nm /280nm absorbance profiles indicated that the Gag^ΔMA^:Ψ^CES^_2_ assemblies contain both protein and RNA (Fig. S5), it was unclear if each assembly contained a single Ψ^CES^_2_ dimer or if multiple dimers were present. To determine RNA distributions within the Gag^ΔMA^ lattices, negative stain EM images of assemblies generated in the presence of 3’-end gold-labeled Ψ^CES^_2_ RNA were obtained. The RNAs used for these studies were prepared by click-chemistry ligation of alkyne-pCp, and azide-functionalized 5-nm gold nanoparticles to the 3’-termini of Ψ^CES^ RNA (Fig. 2d) (see Materials). For each assembly, several dense puncta attributed to individual gold nanoparticles were readily observed at positions that varied among the different assemblies (Fig. 2e). These findings indicate that each assembly contains several Ψ^CES^_2_ molecules, and that the RNAs are heterogeneously distributed within the assemblies.

We next used single particle cryoEM to assess structural information about the RNA organization in the Gag^ΔMA^:Ψ^CES^_2_ assemblies, using identical sample preparation conditions. Two datasets were collected: a less concentrated dataset, which yielded a well-defined ab initio reconstruction that served as a reference, and a more concentrated dataset, which provided higher particle numbers for refinement (Fig. S6). From these, two maps were obtained. A C6 symmetry reconstruction yielded a high-resolution protein map (3.2 Å), which is consistent with previously described immature Gag hexamer structures 34,35,40,44,45. We observed clear density for IP6 molecules bound at the center of each 6HB (Fig. 3a, Fig. S6c) and crystal structures of the CA domains (PDB IDs 4usn, 5i4t) 35,44 fit well into the protein layer density, confirming a canonical Gag hexameric lattice (Fig. 3 c, d, e).

**Figure 3.**
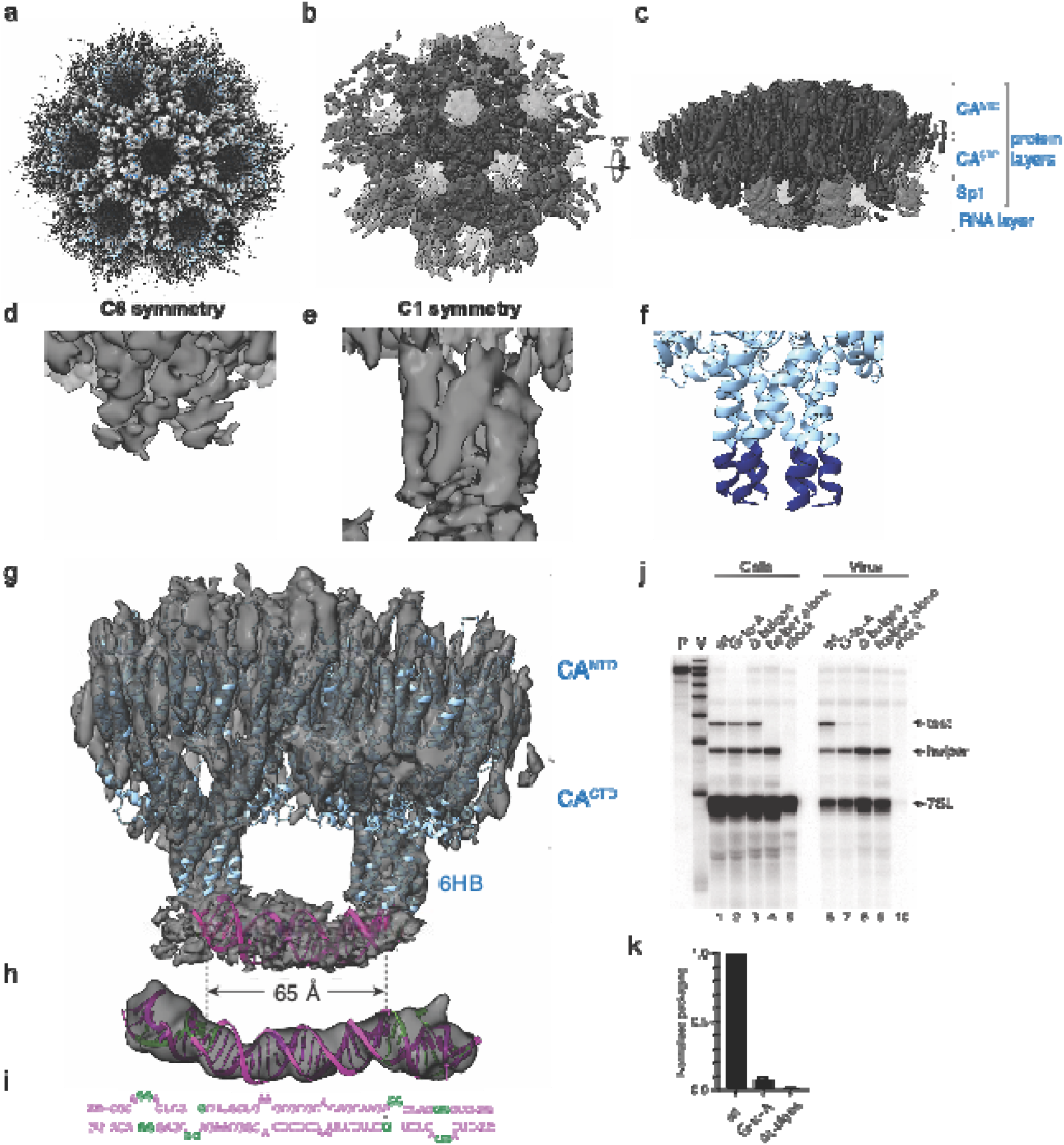
Structural and functional analysis identifies the DIS_2_ as a molecular ruler for Gag hexamer dimerization and reveals a requirement for flanking guanosines on competitive RNA packaging. **a:** CryoEM reconstruction refined on the full immature Gag lattice with C6 symmetry, showing a high-resolution map of the protein in top and side views. PDB entries 4usn and 5i4t were merged and fitted into the protein density (blue). **b, c:** Resolution and density of the CA C-terminal domain is reduced, but density for the 6HB and RNA is enhanced compared to processing with C6 symmetry (Fig. 5). **d:** Zoom in showing the six-helix bundle of the C6 reconstruction with density observed through residue V370. **e**: However, densities for some 6HBs in the non-symmetrized reconstructions extend to the first residue of NC (M378) and connect the protein and RNA layers. **f:** To show the extended SP1 helix, PDB 7ash is shown next to the density. Residues after the mutated T371 (commonly known as the 8^th^ residue in SP1 (T8I) are shown in navy. **g**: A minimal nucleation unit consisting of two dimerized hexamers bound to RNA. These features are consistent with the reconstructions obtained by subtomogram averaging. A structure of DIS_2_ (PDB: 6bg9) was truncated at the exposed guanosines and fitted into the tubular density beneath the lattice (magenta). **h:** The full structure fitted into their reconstructed map (EMDB: 7079), including the exposed guanosines flanking DIS_2_ (green), (**i**) together with a nucleotide representation are shown underneath. **j:** RNase protection assay of samples obtained from co-expression of test vectors with a NL4-3 Ψ^+^ helper. P, undigested probe; M, RNA sizes marker. Lanes 1, 2, and 3: HIV-1 NL4-3 Ψ^+^ helper co-expressed with test vectors containing intact leader (lane 1), G-to-A mutation (lane 2) or Δ bulges mutation (lane 3). Lane 4: HIV-1 NL4-3 Ψ^+^ helper expressed without test RNA. Lane 5: mock transfected cells. Samples obtained from transfected cells (Cells) or viral containing media (Virus) are indicated. Bands corresponding to host 7SL RNA, HIV-1 NL4-3 helper RNA (helper) and co-packaged test RNAs (test) are labeled. **k:** Quantification of packaging efficiency of HIV-1 test vectors under competitive conditions. Packaging efficiency was calculated by dividing the ratio of test vector RNA to helper RNA in the virion sample by the ratio of test RNA to helper RNA in the cells as determined by RNase protection assay and quantified by phosphorimager analysis. For all quantifications, the results represent data from at least three independent transfection experiments.

Refinements of the same set of particles without imposition of symmetry restraints (C1 reconstruction) afforded maps with extended 6HB densities that connect the C-terminus of CA to the NC domain, along with additional tubular density that bridges two neighboring 6HBs (Fig. 3b, c, g). The protein lattice was less well resolved in the C1 reconstructions, and local-resolution analysis revealed that the bridging density was substantially less well resolved than that of the surrounding CA lattice (Fig. S7 a, b). Particle orientation analysis further revealed nonuniform angular sampling and pronounced directional resolution anisotropy in the C1 reconstruction (Fig. S7 c, d, e, f). Consequently, the density did not permit nucleotide-level modeling or direct sequence assignment. However, considering all the individual helical elements within the Ψ^CES^ NMR structure (U5 /AUG helix, Ψ-stem loop, PBS hairpin, and the DIS_2_), only the DIS_2_ helix is of sufficient length to span the density connecting the two 6HBs 27. Indeed, a previously determined DIS_2_ structure can be readily accommodated within the tubular feature of the cryoEM assembly (Fig. 3g).

Several alternative processing strategies were employed, including repeated 3D classification and heterogeneous refinement, local refinement, and signal subtraction using masks that removed different portions of the protein lattice while retaining the 6HB and RNA-associated regions. None of these approaches identified a particle subset with substantially improved definition of the bridging density (Methods). The limited local resolution therefore appears to reflect heterogeneity of the RNA-associated region that could not be resolved by further classification of the available particle population. Additional, weaker densities that lacked a strong connection to the DIS_2_ density were also observed at higher noise thresholds. These additional densities are tentatively attributed to other regions of the Ψ^CES^_2_ RNA that appear to be heterogeneously oriented relative to the DIS_2_ helix.

Notably, in contrast to the C6-refined map (Fig. 3d) and prior wild-type immature lattice structures 34,44, the C1 map displays continuous density for the full SP1 helix (Fig. 3e). Previously, such density had only been reported for a T8I CA-SP1 mutant 45.

### Cryo-electron tomography supports a heterogenous RNA-bridged architecture

Because assemblies exhibited heterogeneity in both size and RNA distribution, we additionally applied cryo-electron tomography (cryoET) and subtomogram averaging to confirm this structural organization. CryoET data were obtained for Gag^ΔMA^ assemblies with IP6 and both the Ψ^CES^_2_ and DIS_2_ RNAs (Fig S8). Reconstructions from all manually picked particles resolved the three expected layers: CANTD, CACTD (combined as protein layer), and NCSP2p6 bound to RNA (RNA layer) (Fig. S8 and Fig. S9). In these initial reconstructions, density for the RNA layer was more diffuse compared to the two protein layers, indicating that the RNA is not uniformly aligned with the CA domains. Layer-focused local refinements (Fig. S8) revealed that the protein and RNA layers are not translationally shifted but exhibit rotational heterogeneity (Fig. S11). Classification of particles displaying strong density in the RNA layer spanning two adjacent 6HBs yielded reconstructions from 767 particles for the Ψ^CES^_2_ dataset and 657 particles for the DIS_2_ dataset. These reconstructions, both without imposed symmetry, closely resembled those obtained by cryoEM, with tubular densities bridging two adjacent 6HBs (Fig. S9 and Fig. S10). Further denoising and processing using IsoNet2 did not improve the definition of this feature (Methods). Although neither of the subtomogram averages exhibited resolution sufficient to identify structural features expected for an RNA helix, the similarity of the independent cryo-ET datasets obtained for assemblies prepared with the Ψ^CES^_2_ and DIS_2_ RNAs, and by cryoEM for the Ψ^CES^_2_ containing assemblies, support our conclusion that the tubular density bridging the adjacent 6HBs is due to the DIS_2_ helix.

Whereas the CA lattice is highly ordered in both the single-particle and tomographic datasets, RNA density outside the prominent inter-hexamer feature is diffuse and variably positioned. Local-resolution analysis further revealed substantially lower resolution in the RNA associated region compared with the surrounding CA lattice. In addition, directional FSC analysis identified pronounced resolution anisotropy in the C1 reconstruction, suggesting that nonuniform angular sampling may also contribute to the limited definition of the RNA-associated density (Fig. S7). Extensive 3D classification, focused refinement, and signal-subtraction approaches did not isolate a particle population with substantially improved RNA density. Similarly, denoising of the tomographic data did not improve the definition of the RNA-associated density. The RNA layer therefore does not appear to adopt a single rigid arrangement relative to the extending CA lattice. This heterogeneity could reflect multiple RNA-binding configurations or different stages of lattice growth. One possibility is that a comparatively constrained RNA architecture contributes to formation of an initial Gag assembly intermediate, whereas additional Gag molecules interact more flexibly with the surrounding RNA as the lattice expands. This model could reconcile the defined two-hexamer architecture observed in the reconstructions with the presence of multiple, heterogeneously distributed Ψ^CES^_2_ molecules in the larger assemblies.

### DIS_2_ guanosine clusters promote RNA packaging

In the structural model described above, placement of the DIS_2_ within the bridging density positions clusters of non-base-paired or non-canonically paired guanosines flanking DIS_2_ beneath the 6HBs and proximal to the C-terminal NC domains (Fig. 3g, h, j) 27,46. Since HIV-1 NC binds with high affinity to oligonucleotides that contain exposed guanosines 14,47,48, this arrangement suggests that the DIS_2_-flanking guanosines could contribute to Gag-RNA interactions during genome selection. The influence of these DIS_2_ guanosines on RNA encapsidation was evaluated using a competitive *in situ* RNA packaging assay 14,27 using RNAs containing either G-to-A substitutions (G240A, G241A, G247A, G272A, G273A, G278A, and G279A) or mutations that eliminate non-canonical base pairing (deleted: A239-A242, G247, A271-G273, G278-G279). Free energy calculations indicate that these substitutions should not alter the secondary structure of the RNA 49. Human embryonic kidney 293T cells were co-transfected with plasmids that produce vector RNAs containing the wild type (Ψ^+^, which also encodes for viral proteins) and mutant (Test) leader sequences. When co-expressed at similar levels, Ψ^+^ and Test vector RNAs with native leader sequences were packaged into HIV-1 virus-like particles with similar efficiencies (Fig. 3j, k). In contrast, significant packaging defects were observed for both the G-to-A mutated Ψ^CES^ RNA (7.5% ± 2.2%), and the Ψ^CES^ RNA lacking non-canonical base pairs (1.4% ± 0.08%). Thus, the exposed guanosine residues flanking the DIS_2_ helix are required for efficient competitive genome packaging.

## CONCLUSIONS

Our studies reveal that the dimeric form of the HIV-1 RNA packaging signal (Ψ^CES^_2_) promotes in vitro assembly of 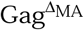 molecules whereas the monomeric Ψ^CESm^ analog does not. Assembly also requires IP6 and can be promoted by the dimeric DIS_2_ region of Ψ^CES^_2_. In the presence of either Ψ^CES^_2_ or DIS_2_, the CA domains assemble into lattices similar to those observed previously for immature virus-like particles. Previous studies proposed that Gag dimers seed assembly by formation of incomplete hexamers at lattice edges 50. Our findings place DIS_2_ at the center of this process, providing both specificity through RNA dimerization and structural continuity between neighboring hexamers. The requirement of IP6 for driving cooperative binding to Ψ^CES^ suggests that its role extends beyond stabilization of the SP1 6HB 33,40,51 to include modulation of NC-RNA interactions. The appearance of a lower-molecular weight intermediate preceding higher-order assembly indicates that initial organization occurs on the RNA scaffold before lattice growth proceeds. This interpretation is consistent with earlier studies showing that RNA dimerization, rather than RNA abundance alone, is critical for packaging specificity 52. Notably, assemblies formed with the wild-type CA sequence exhibit continuous density for the SP1 helices, even in the absence of mutations previously shown to stabilize the 6HB 45,53.

The requirement of a dimeric DIS_2_ for efficient assembly is consistent with proposals that the dimeric packaging signal is an architectural factor rather than a passive cargo 22,54. In its dimeric state, DIS_2_ forms a helix sandwiched by clusters of non-base paired or non-canonically base paired guanosines, a feature conserved among lentiviruses 55. Separation between the guanosine clusters (∼ 65 Å) matches the separation between 6HB structures in CA-SP1-containing assemblies 34, enabling DIS_2_ to template cooperative IP6-dependent formation of two adjacent Gag hexamers. Extension of the 6HBs to the SP1-NC boundary places the downstream NC domains in close proximity to the clusters of exposed guanosines (known targets of NC’s zinc knuckles 27,46,48), and identifies an RNA recognition function for SP1, an established antiviral target 56-58. RNA mutations designed to eliminate NC binding by replacing the guanosines by adenosines, or by removing them altogether, severely impair RNA packaging into virus-like particles. A caveat of the present studies is that we were unable to obtain resolution by cryoEM or cryoTM sufficient to identify residues or observe helical structure in the tubular densities attributed to the DIS_2_ helix. However, multiple lines of evidence support this assignment, including: (1) the assemblies are only obtained for complexes with the dimeric Ψ^CES^_2_ (but not the monomeric Ψ^CESm^ ); (2) similar tubular densities were observed by both cryoEM and cryoET for assemblies containing Ψ^CES^_2_ and DIS_2_-only RNAs, and (3) based on NMR and cryoET studies of RNA fragments, DIS_2_ is the only helix of sufficient length to span the two 6HBs.

In summary, our findings suggest that DIS_2_ functions as a molecular ruler for organizing Gag hexamers into a geometry optimized for Gag lattice expansion 50 and identifies DIS_2_ as a central determinant that links RNA dimerization with NC binding specificity, Gag nucleation, and CA lattice propagation.

## ACKNOWLEDGMENTS

We thank Yu Chen, and staff at the Howard Hughes Medical Institute (HHMI), and staff at the University of Maryland, Baltimore County (UMBC) for technical assistance and helpful suggestions. We also thank Xiaowei Zhao, Rui Yan, Zhiheng Yu, James Jung, Adamo Mancino, Nicholas Spellmon, and Shixin Yang at the HHMI Janelia CryoEM facility for help in microscope operation and data collection.

## AUTHOR CONTRIBUTIONS

N.M.H., S.K., A.B., C.P., A.L., F.A., A.A., K.B., S.A.G., A.T., B.K.G., O.P., and M.F.S. designed research; N.M.H., S.K., A.B., C.P., A.L., F.A., A.A., K.B., S.A.G. and C.B. performed research; N.M.H, B.K.G., and O.P. contributed new analytical tools; N.M.H., S.K., B.K.G., O.P., and M.F.S. analyzed data; and N.M.H., B.K.G., O.P., and M.F.S. wrote the paper.

## DATA AND CODE AVAILABILITY

Maps were deposited at the EMDB with accession numbers EMD-76853, EMD-76854, EMD-76863, EMD-76864.

