## Supplementary Materials and Figures for "Structural Basis for Dimeric Genome Selection by HIV-1"

#### **Methods**

#### **Plasmids**

*E. coli* DH5 α (*fhuA2 lac(del)U169 phoA glnV44 Φ80’ lacZ(del)M15 gyrA96 recA1 relA1 endA1 thi-1 hsdR17*) was used to generate plasmids that were cloned in this study. The cells were grown in LB medium at 37°C and harvested after overnight cultures. The QIAprep Spin Miniprep Kit was used for plasmid purification.

The plasmids for the expression of HIV-1 Gag polyprotein CANCSP2p6 (V128-Q500) and all other tested constructs (Fig. S1) were derived either from pETM11 (derived from pBR322; G. Stier) or pLATE11 using the restriction free cloning approach. Point mutations were inserted by site directed mutagenesis ^1^.

#### **Protein expression and purification**

*E. coli* BL21 (DE3) cells (*E. coli B dcm ompT hsd(r_B_^-^m_B_^-^)gal*) were used to express the various recombinant proteins. The cells were grown at 37°C in LB medium and induced with 1 mM IPTG at an OD_600_ of 0.8, followed by overnight incubation at 17°C.

After protein expression, harvested cells were resuspended in 20 mM Mes (pH 6.0), 500 mM NaCl, 1 M Urea, 100 μM ZnCl_2_, 1 mM PMSF, 2 mM TCEP, and 1x protease inhibitor cocktail (Fisher) and lysed using a microfluidizer. 1.5% (v/v) PEI (10% stock solution) was added dropwise to the cleared lysate and stirred for 1 hour at 4°C. After an additional centrifugation step, ammonium sulfate was added to 50% and stirred for another 2 hours at 4°C to separate residual nucleic acids from the proteins. The resulting protein pellet was resuspended in 20 mM Mes (pH 6.0), 300 mM NaCl, 1 mM PMSF, and 2 mM TCEP, and dialyzed overnight at 4°C using a 10 kDa molecular weight cut-off membrane in the same buffer. The next day, the protein solution was applied to a 5 ml FF GST column (Cytiva). After washing with 20 column volumes (CV) of dialysis buffer, the protein was eluted with a 10 CV gradient of 20 mM reduced glutathione. The eluted protein was then cleaved with TEV-protease and dialyzed overnight at 4°C against 20 mM Mops (pH 7.4), 300 mM NaCl, 1 mM PMSF and 2 mM TCEP. The cleaved protein was passed over a second GST column to remove the tag and then concentrated using an Amicon Ultra centrifugal filter (Merck-Millipore) with a 10kDa cut-off. Final purification and buffer exchanged were performed via size-exclusion chromatography using a Superdex 75 column (Cytiva) and 20 mM Mops (pH 7.4), 140 mM KCl, 10 mM NaCl, and 5 mM TCEP. The purified protein was concentrated to approximately 100 μM and flash-frozen in small aliquots to prevent freeze-thaw cycles.

Proteins tested without an affinity tag were purified using an SP ion exchange column (16/60; Cytiva), eluting with a buffer containing 1M NaCl.

#### **RNA in vitro transcription and purification**

DNA templates for Ψ^CES^_2_ and Ψ^CES^_mon_ were generated by PCR amplification (EconoTag PLUS 2x Master Mix, Lucigen) from a pUC57 plasmid containing the desired sequences, preceded by the Top17 promoter sequence (5′-TAATACGACTCACTATA-3′) for RNA transcription. Reverse amplification primers had their first two 5’-residues 2’-O-methylated to reduce self-templated run-on transcription ^2^. DNA templates were purified via sodium acetate and ethanol precipitation prior to the transcription reaction.

DNA for the shorter DIS construct was purchased form IDT and included the same 5’ modifications and the T7 primer.

Large scale in vitro transcription using T7 RNA polymerase was performed as described previously^3^. Reactions contained approximately 1 mg of the DNA template, 20 mM MgCl_2_, 3 mM NTPs (Cayman), 2 mM spermidine, 5 mM DTT, 20% (v/v) DMSO, 0.1% (v/v) Triton X-100, 40 mM Tris-HCl (pH 9.0), and varying amounts T7 RNA polymerase (0.5-1 mg). Reactions were incubated overnight at 37°C and then quenched with 500 mM EDTA solution (pH 8.0).

After quenching, samples were denatured at 100°C for 5 minutes, snap cooled on ice and mixed with 2x RNA loading dye. The reactions were purified on a 7.5 M urea polyacrylamide gel (19:1 acrylamide/bisacrylamide; SequaGel; National Diagnostics) run at a constant power of 20 W for 18 hours. RNA was visualized by UV shadowing and recovered using the crush-and-soak method. Briefly, gel slices were broken into small pieces and incubated overnight at 37°C on a rocking shaker with 1x TBE (44.5 mM Tris-Boric Acid (pH 7.5), 0.2 mM MgCl_2_, and 2 mM EDTA). Gel debris was removed using a 0.45 μm pore-size filter, and RNAs were purified via sodium acetate and ethanol precipitation.

#### **Gold labeling of RNA**

Gold-labelled RNA was generated in two steps. First, pCp-alkyne (Jena Biosciences) was ligated to the 3’ end of ψ_CES_ RNA using T4 RNA ligase (Thermo Fisher). Nine nanomoles of RNA were used, and the reaction was incubated overnight at 4°C following the manufacturer’s protocol. The RNA was then purified using the Monarch Spin RNA Cleanup Kit (NEB) according to supplied instructions.

In the second step, an azide functionalized 5 nm gold nanoparticle (Nanopartz) was conjugated to the alkyne-modified RNA via copper-catalyzed azide-alkyne cycloaddition (CuAAC) click chemistry, using the THPTA-based CuAAC Biomolecules Reaction Buffer Kit (Jena Biosciences). To obtain sufficient material, nine individual reactions were performed in parallel. After 1 h incubation at 37°C, the reactions were quenched, and the labeled RNA was gel-purified as described above.

#### **Electrophoretic mobility-shift assays (EMSA)**

RNA samples (2 μM) were prepared in water, denatured at 100°C for 2 min, and then incubated overnight at 37°C in 20 mM Mops (pH 7.4), 140 mM KCl, 10 mM NaCl, ±1 mM IP_6_, and 5 mM TCEP (PI buffer) to reduce RNA oligomerization and facilitate proper dimerization. Protein samples were diluted in PI buffer to their respective concentrations and incubated on ice for 20 min. Protein and RNA were then mixed at indicated ratios and incubated for 20 min at room temperature.

Following incubation 20 % v/v 50% glycerol was added. Samples containing 1 μM RNA were loaded onto 1% agarose gels pre-stained with ethidium bromide. Electrophoresis was performed in pre-chilled 1x TB buffer (44.5 mM Tris-Boric Acid (pH 7.5), 0.2 mM MgCl_2_) until adequate band separation was achieved, as assessed by migration of bromphenol blue and xylene cyanol in a control lane.

#### **Analytical size exclusion chromatography**

RNA, protein, and complex samples were prepared as described for the EMSAs. A 200 μl sample volume was prepared at a 1 μM RNA concentration and injected on a 10/300 Superdex 200 increase column (Cytiva). The samples were run at 4°C on an Äkta pure system, with absorbance monitored at 260 nm, 280 nm, and 215 nm.

#### **Negative staining and transmission electron microscopy**

RNA, protein, and complex samples were prepared as described for the EMSAs. After incubation the complex was diluted to a final concentration of 0.1 μM just before application onto the grid.

Carbon-supported film copper grids (200 mesh) were glow-discharged (GloQube Plus) immediately prior to sample application. A 4 μl aliquot of the diluted complex was applied to the grid and incubated for 1 minute. Excess liquid was quickly blotted using filter paper, followed by two rapid washes in H_2_O (brief incubation and fast blotting between washes). The grid was then stained in 0.75% (w/v) uranyl formate for 1 minute, after which excess stain was blotted away. Grids were dried for at least 4 hours before insertion into the microscope.

Imaging was performed using a Hitachi HT7800 transmission electron microscope (TEM) operating at 120 kV, equipped with a CMOS AMT Nanosprint15 B camera.

#### **Single particle Cryogenic electron microscopy data acquisition and analysis**

RNA, protein, and complex samples were prepared as described for the EMSAs and immediately applied (4 μl) to glow-discharged Quantifoil R1.2/1.3 300-mesh Au grids. Grids were plunge frozen using a Vitrobot Mark IV (ThermoFisher) with a blot force of 4 and a blot time of 2 s.

Single particle cryo-EM data were collected at the Cryo-EM facility at the HHMI Janelia Research Campus. Two datasets were acquired on Titan Krios microscopes (Thermo Fisher Scientific) operated at 300 kV. The microscope used for dataset 1 was equipped with a high-brightness field-emission gun (X-FEG), a spherical aberration corrector, and a Gatan K3 direct electron detector. The microscope used for dataset 2 was equipped with a cold field-emission gun, a Thermo Scientific Selectris X energy filter, and a Thermo Scientific Falcon 4i direct electron detector. Data were collected using SerialEM at a pixel size of 0.426 Å in super-resolution counting mode for dataset 1 and 0.94 Å for dataset 2, with a total exposure of 50 electrons/Å² fractionated over 40 frames and a target defocus range of −1.2 to −2.0 µm.

Image processing and map calculations were performed in cryoSPARC (v4.6.0) ^4^. Raw movies were corrected for beam-induced motion using MotionCor2, and CTF estimation was performed with CTFFIND4 ^5^, as implemented in cryoSPARC. Particles were identified using a combination of blob-picking, filament tracing, and the template pickeing using EMD4016^6^ as a template. Particles were extracted in 880-pixel boxes and Fourier cropped to 440 pixels (Fig. S6a).

For dataset 1, 16,423 movies yielded 81,695 particles after iterative reference free 2D classification, removal of poorly defined classes, merging of particles obtained using the different picking strategies, and removal of duplicate particles. The selected particles were used for ab initio reconstruction without imposed symmetry (C1), followed by homogenous refinement. The resulting reconstruction was used as an initial reference for processing dataset 2.

Dataset 2, a more concentrated sample, comprised 3,171 micrographs and yielded 164,683 particles after particle picking, iterative 2D classification, particle selection, and duplicate removal. Following initial homogenous refinement against the reconstruction obtained form dataset 1, the amp was recentered by volume alignment on two adjacent 6HBs associated with the strongest intervening RNA density. Particles were reassigned and re-extracted at the recentered coordinates, and duplicate particles were removed, yielding a final set of 148,351 particles. Final homogeneous refinements were performed without imposed symmetry (C1) to preserve asymmetric RNA-associated density and with C6 symmetry to optimize features of the locally hexameric Gag lattice. Gold-standard Fourier shell correlation (FSC) estimates using the 0.143 criterion indicated global resolutions of 3.8 Å for the C1 reconstruction and 3.2 Å for the C6 reconstruction.

To determine whether conformational or compositional heterogeneity limited resolution of the RNA-associated density, we extensively explored alternative particle classification and refinement strategies. These included repeated ab initio reconstructions with different numbers of classes, heterogeneous refinement, 3D classification with two to five classes, non-uniform refinement, and local refinement using masks encompassing different regions of the Gag–RNA assembly. Particle subsets identified by these approaches were additionally subjected to further 2D classification and homogeneous refinement. Although these procedures produced differences in the quality of the protein lattice, none identified a particle subset in which the density connecting adjacent 6HBs was substantially better resolved than in the consensus C1 reconstruction.

We additionally tested signal-subtraction and focused-refinement strategies designed specifically to improve the RNA-associated density. Protein density was subtracted from individual particles using masks encompassing different portions of the Gag lattice, including masks that removed the majority of the protein layer while retaining either the complete 6HB region or more restricted subsets of the 6HB density. The resulting signal-subtracted particle sets were subjected to focused local refinement and additional classification. These approaches did not improve the continuity or resolution of the density connecting adjacent 6HBs. The consensus C1 reconstruction was therefore used for interpretation of the asymmetric RNA-associated density, whereas the C6 reconstruction was used for analysis of the higher-resolution Gag lattice and IP6-binding site.

#### **Single particle tomography data acquisition and analysis**

RNA, protein, and complex samples were prepared as described for the EMSAs. After addition of 10nm BSA Gold tracer, 4 μl were immediately applied to glow discharged C-Flat 2/1 300-mesh Cu grids grids and frozen using a EM PG2 Automated Plunge Freezer (Leica) (5 sec blot time).

Cryotomograms were collected at the University of Utah on a Krios G2 (ThermoFisher) operated at 300 kV, equipped with a Gatan K3 camera. Four tomograms were collected at a magnification of 64 000 x, which corresponds to a pixel size of 1.407 Å using Serial EM in counting mode, with a total dose of 150 electrons/Å^2^ over 61 frames and an angular range of -60° to +60°, an angular increment of 2° starting from 0°, and a defocus of -4 to -5 μm.

Tilt series were aligned using IMOD^7^ and sub tomogram averaging was carried out using the Dynamo software package ^8^. Sub volumes were extracted from 2x or 4x binned data in box sizes of 64 or 32 px (Fig. S7) after assemblies were manually picked.

Picked particles were initially aligned imposing C6 symmetry. The specific averaging parameters for each run are summarized in Table S1. After each alignment round, the resulting volumes were examined for the presence of the expected capsid lattice and the targeted structural feature. Following the initial alignment, a multi-reference alignment was performed using (i) the map from the initial run and (ii) a copy of the map with the RNA density removed. The mask for this run was generated in Dynamo using the dmask tool, creating a cylinder with a radius of 30 px, soft-padded with a Gaussian filter of 4 ox, and positioned to cover the RNA layer.

To maximize recovery of particles, we performed a 6-fold symmetry expansion. Magnesium atoms were placed in the center of the central CA hexamer and in the centers of the six adjacent CA hexamers. Using the transformation matrix for each particle (calculated with dtplace), a new table was generated in which each particle was transformed seven times according to the Mg atom positions. Duplicate particles were removed in a subsequent alignment run.

Particles were then re-extracted around a single six-helix bundle (6HB) using the same Mg-based positioning strategy, followed by multi-reference alignment with a sphere-shaped mask (radius 5 px, Gaussian filter 2 px) centered on the 6HB. Volumes were screened for the presence of a 6HB with connected RNA density. Selected particles were subsequently re-extracted around the adjacent 6HB and screened again. Finally, particles were re-extracted from the midpoint between these two 6HBs for the final alignment.

To assess structural heterogeneity between the protein and RNA layers, local refinements were performed using masks covering either the protein or the RNA. The resulting x, y, and z positions as well as Euler angles were compared (Fig. S10). Fourier shell correlations (FSCs) between even/odd maps were calculated inside soft-edged Gaussianmasks splitting particles between different tomograms randomly into half using the dfsc subroutine in Dynamo. The gold-standard resolution estimates (0.143 cutoff) were used to assess the resolution.

To assess whether denoising could improve visualization and alignment of the comparatively weak RNA-associated density, tomograms were additionally processed using IsoNet2. Denoising was performed using independently reconstructed even- and odd-frame tomograms to minimize propagation of correlated noise during neural-network training and reconstruction. IsoNet2 processing substantially improved the visual contrast of the tomograms but did not yield improved subtomogram averages or enhance the definition of the tubular density connecting adjacent 6HBs. The original, non-denoised tomograms were therefore retained for the quantitative subtomogram averaging and FSC analyses reported here.

#### **Structural analysis and visualization**

Map examination, PDB model fitting, and figure rendering were performed with ChimeraX^9^. Data were plotted using Python.

#### **HIV-1 NL4-3 based helper and test vector plasmids.**

HIV-1 NL4-3 test vector with intact 688-base 5′ leader and HIV-1 NL4-3 Ψ+ helper encoding HIV-1 gag, gag-pol, tat and rev proteins has been described previously ^10^. To create HIV-1 NL4-3 test vectors containing mutations in the leader, synthetic DNA fragments with desired changes in the sequence were ordered from Twist Bioscience HQ ( South San Francisco, CA). Synthetic DNA fragments were directly cloned into HIV-1 NL4-3 test vector, replacing the intact leader sequence. GsIIAs mutant contains Gs substituted with As at positions 240, 241, 247, 272, 273, 278 and 279. Δ bulges mutant contains deletions of the bases 239-242, 247, 271-273 and 278-279.

#### **Viral RNA production and RNA analysis.**

Co-transfection of 293T cells with helper and vector plasmids and RNA isolation from virus and cells were performed as described previously ^10^. Plasmid pSKh136b used as template to generate riboprobe was previously described ^11^*.* Generated riboprobe contains fragments of 7SL RNA and of an SV40 promoter with 5’ adjacent sequence of the HIV-1 NL4-3 present in the vectors used in this study. Due to difference in SV40 flanking sequences, this riboprobe protected fragments of differing sizes for helper and test RNAs (143 and 190nt correspondingly). RNase protection assays performed as previously described ^11^. Products of the RNase protection assays were quantified by phosphorimaging using standard methods.

### **Supplemental Figures**

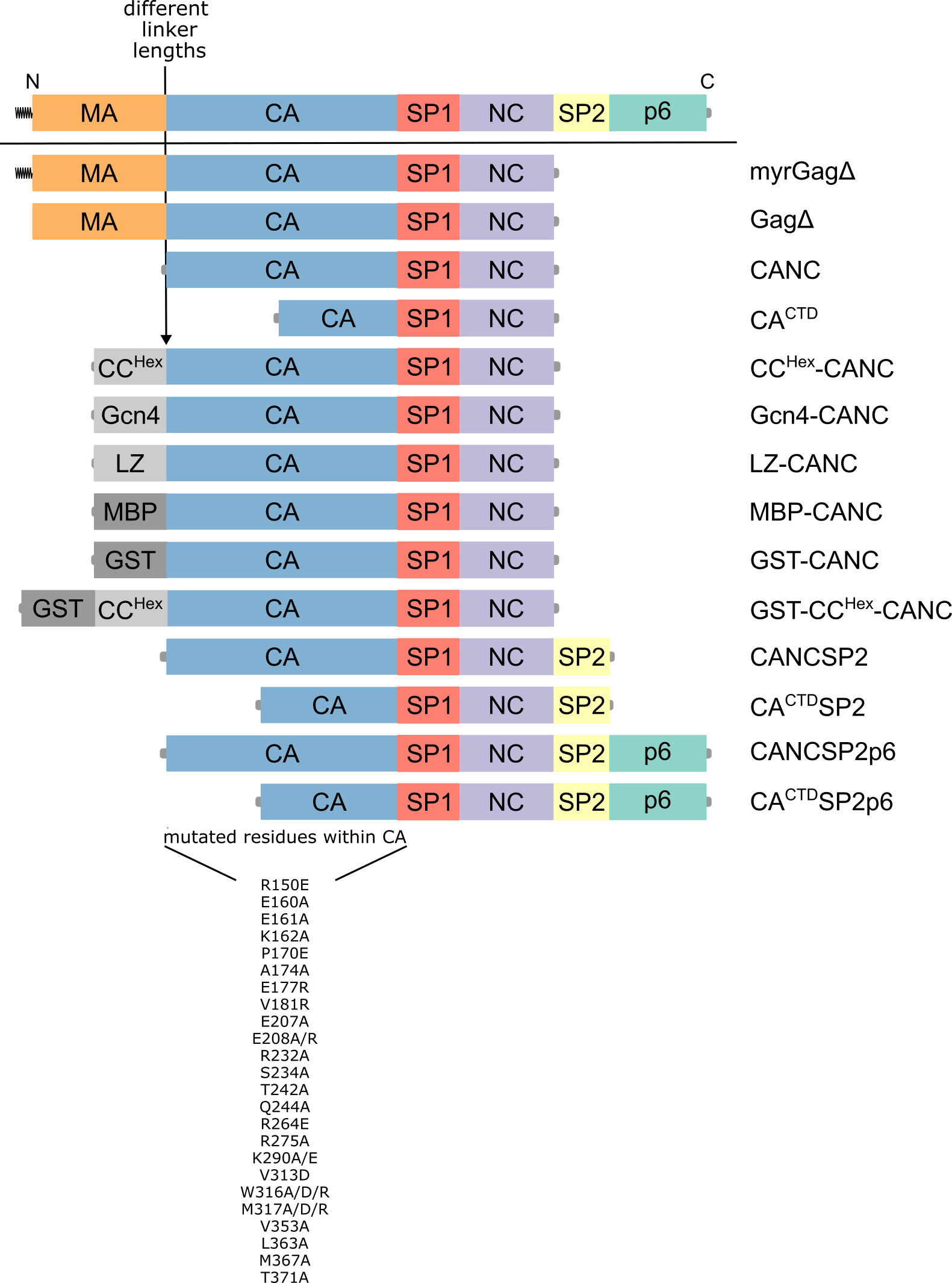

**Figure S1: Schematic representation of tested protein constructs.** Multiple combinations of CA-domain mutations were assessed, none of which enhanced solubility upon formation if protein-RNA complexes.

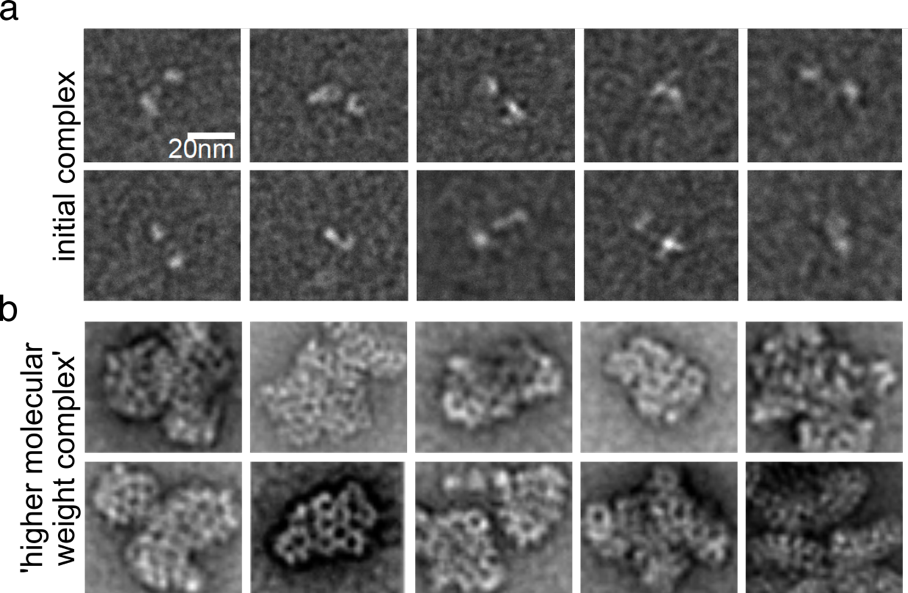

**Figure S2: Negative stain electron microscopy of Gag^ΔMA^:Ψ^CES^_2_ complexes. a:** The initial complex shows a dumbbell-shaped structure. The higher molecular weight assemblies exhibit the well-studied Gag hexameric lattice formation. Scale bar = 20 nm.

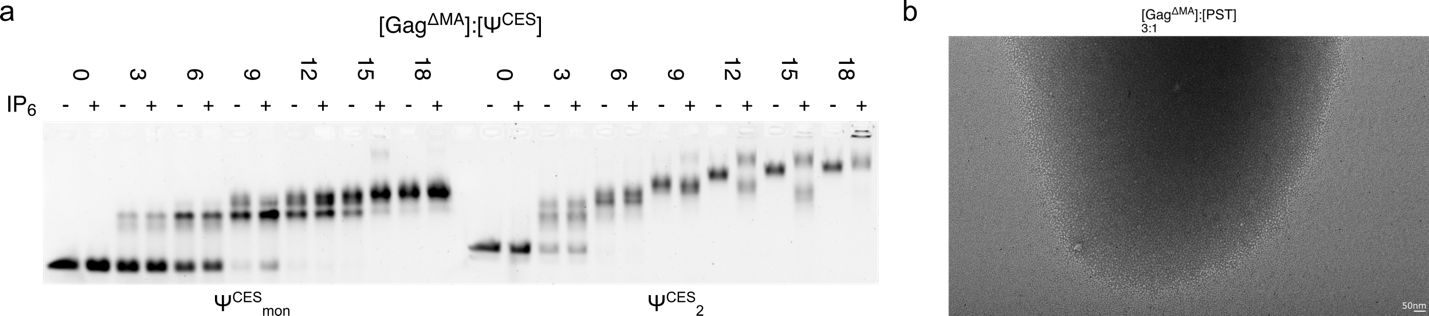

**Figure S3: A system to stimulate cellular selective packaging of Ψ^CES^_2_ by Gag in vitro.** **a:** A gel shift assay comparing complex formation of Ψ^CESm^ (left) and Ψ^CES^_2_ (right). Only Ψ^CES^_2_ supports the formation of higher molecular weight assemblies in the presence of IP_6_. Increasing Gag: Ψ^CES^ ratios from 1:0 up to 1:18 are shown in presence and absence of IP_6_ (1 mM). b: Negative stain image showing that nder the same experimental conditions as the DIS_2_ RNA (3:1 protein:RNA ratio), the Ψ-stem loop fails to form higher-molecular-weight complexes. Scale bar = 50 nm.

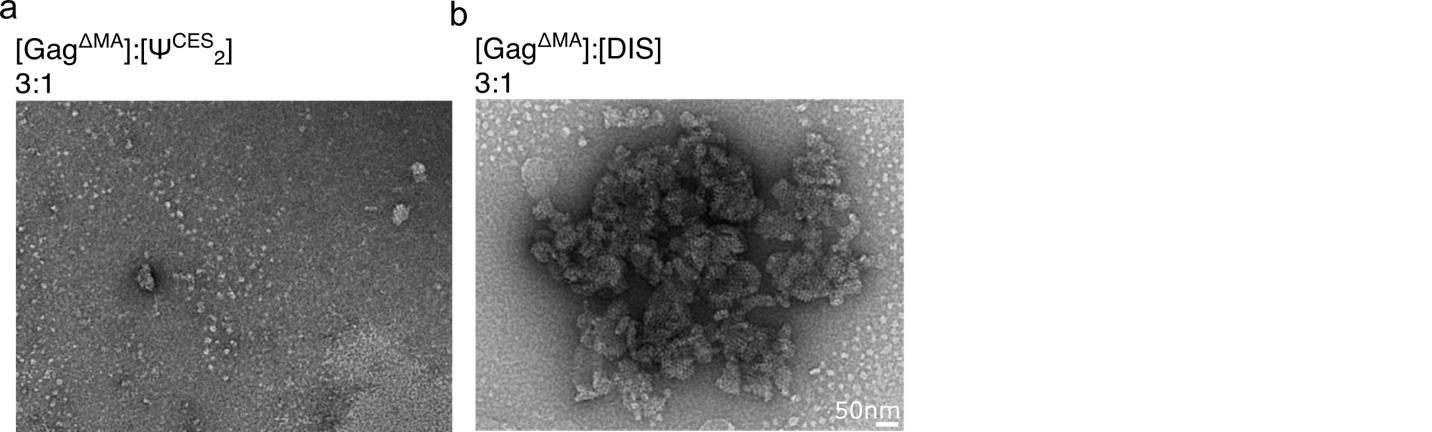

**Figure S4: The DIS_2_ alone is more efficient to facilitate assembly of the hexameric capsid lattice.** Negative-stain EM at a 1:3 ratio reveals minimal hexameric lattice formation for Ψ^CES^_2_, (**a**) while numerous virion-like assemblies are seen for DIS_2_ (**b**). Scale bar = 50 nm.

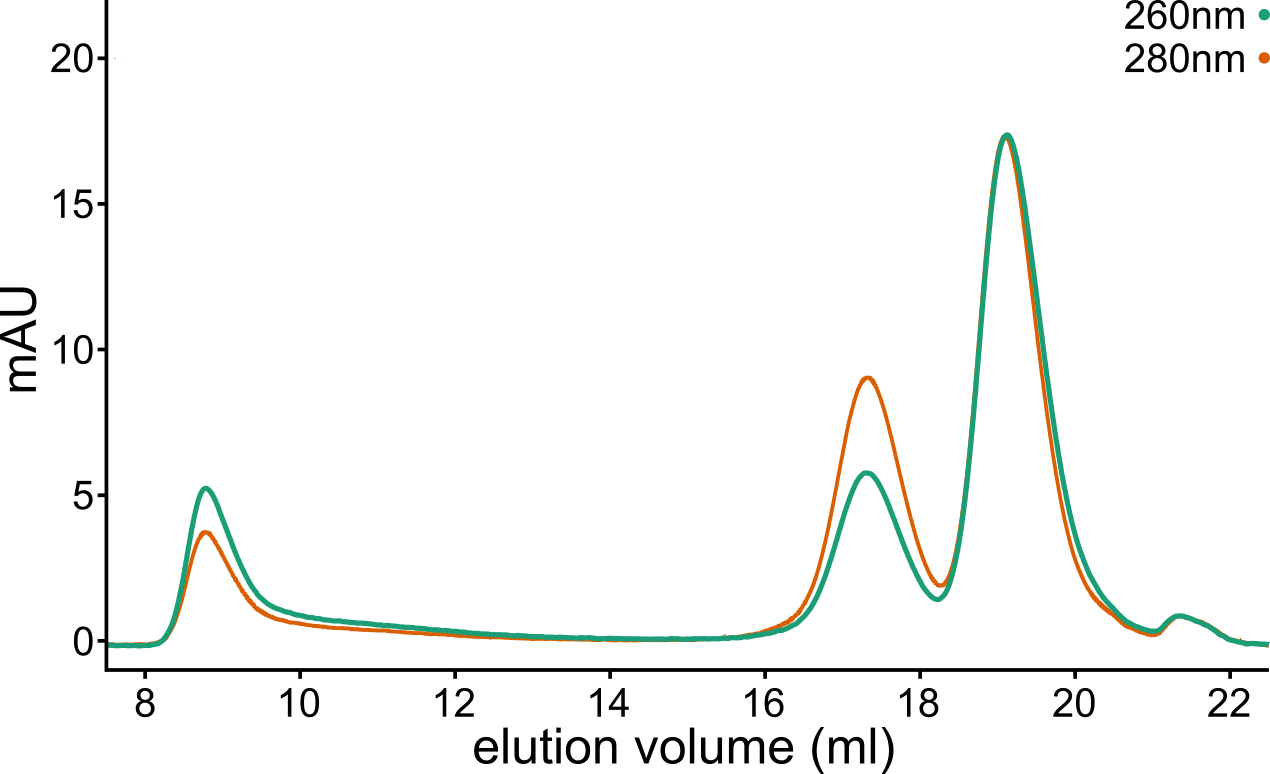

**Figure S5: Assessing RNA recruitment into the hexameric capsid lattice.** The presence of RNA in the higher molecular weight assemblies is confirmed by analytical SEC, with a higher 260 nm absorbance (green) compared to 280 nm (orange). The protein:RNA ratio is 1:12.

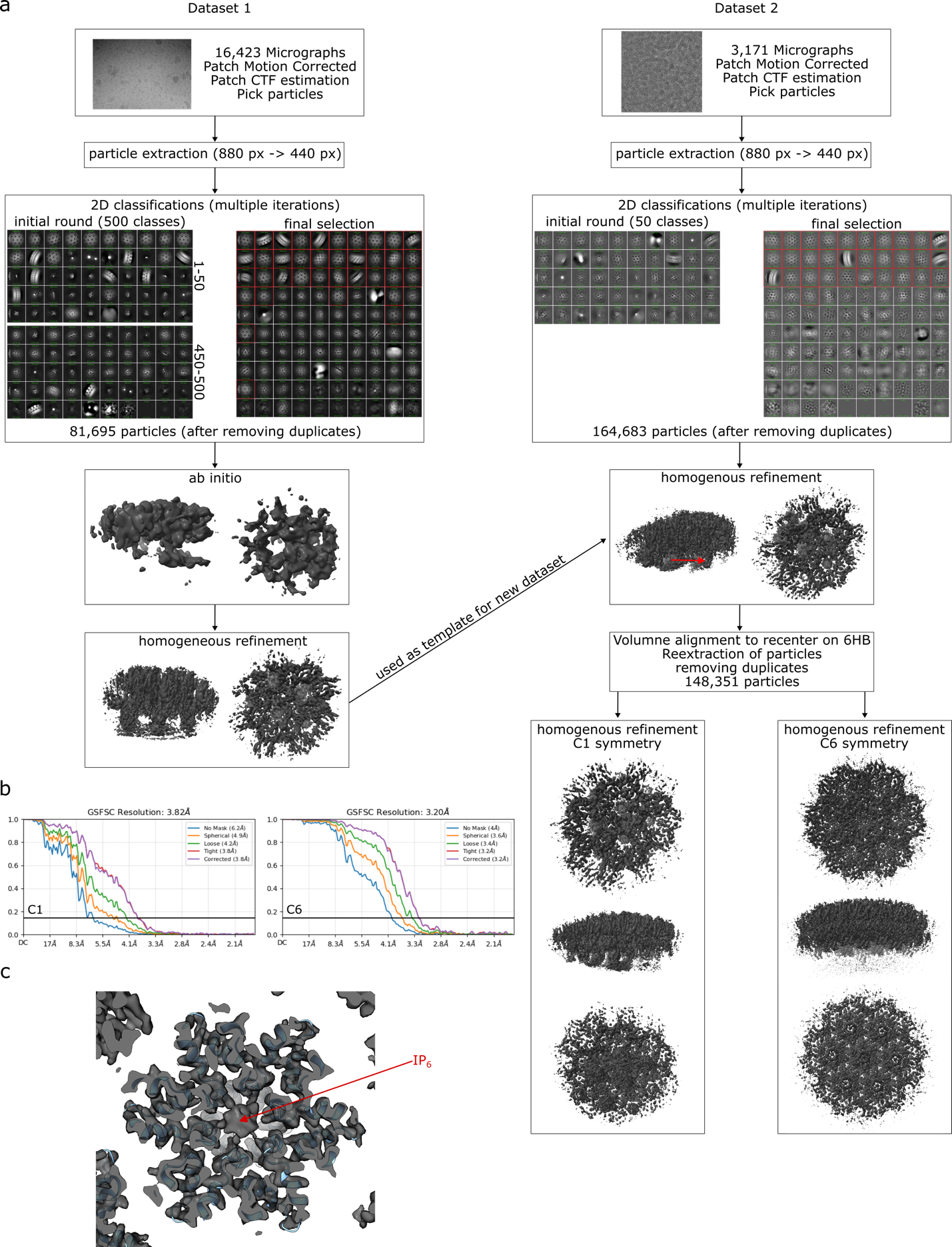

**Figure S6: Workflow and results of single-particle cryo-EM. a:** Flow diagram outlining processing of the two collected datasets. After patch motion correction, CTF estimation, particle picking, extraction, and iterative 2D classification, 16,423 micrographs yielded 81,695 particles (left). These particles were used for ab initio map generation in C1 symmetry, followed by homogenous refinement. The resulting map served as a template for processing the second dataset (right), which after motion correction, CTF estimation, particle picking, and curation yielded 164,683 particles. Homogenous refinement followed by recentering on two 6HBs with the most prominent RNA density, re-extraction, and duplicate removal resulted in 148.351 particles. Final maps were calculated by homogenous refinement with either C1 symmetry (to optimize RNA density) or C6 symmetry (to optimize protein density). **b:** Gold-standard FSC estimates (0.143 cutoff) indicated resolutions of 3.8 Å ( C1) and 3.2 Å (C6). **c:** Density for IP_6_ is readily observed in the higher resolution C6 refined map.

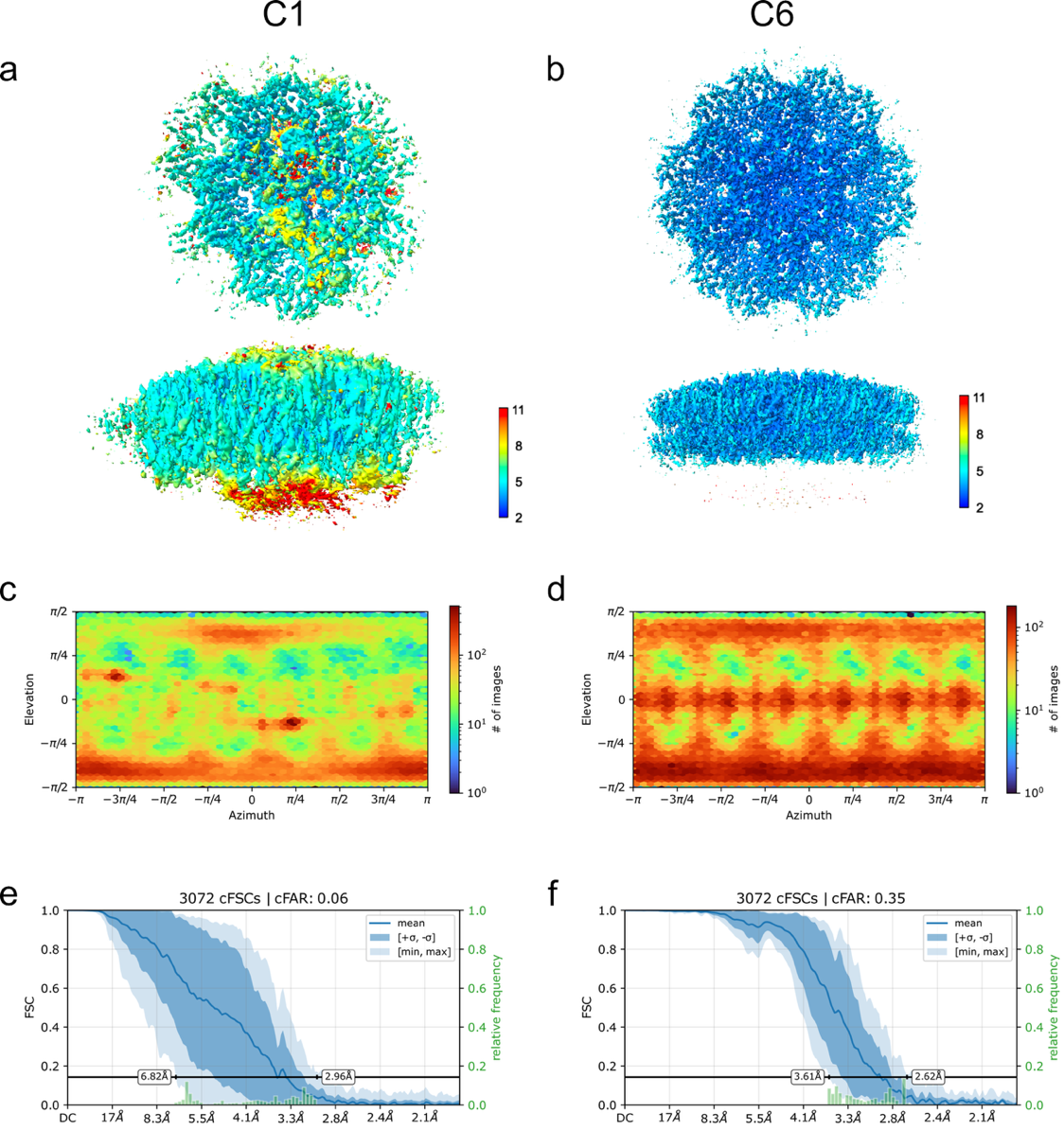

**Figure S7:** **Local-resolution estimates, particle orientation distributions, and directional resolution anisotropy of the C1 and C6 cryo-EM reconstructions.** a/b: Top and side views of the C1 (a) and C6 (b) reconstructions, colored according to local-resolution estimates calculated in CryoSPARC and displayed on the sharpened density maps. Colors indicate estimated local resolutions from 2 to 11 Å. The C1 reconstruction exhibits greater spatial variation in resolution, with comparatively poorly resolved density beneath the ordered CA-SP1 lattice. c/d: Particle viewing-direction distributions for the C1 (c) and C6 (d) reconstructions, revealing nonuniform angular sampling. e/f: Directional resolution analysis based on 3,072 conical Fourier shell correlation (FSC) curves for C1 (e) and C6 (f). Directional resolution estimates ranged from 2.96 to 6.82 Å for C1 (cFAR = 0.06) and from 2.62 to 3.61 Å for C6 (cFAR = 0.35), indicating greater directional resolution anisotropy in the C1 reconstruction.

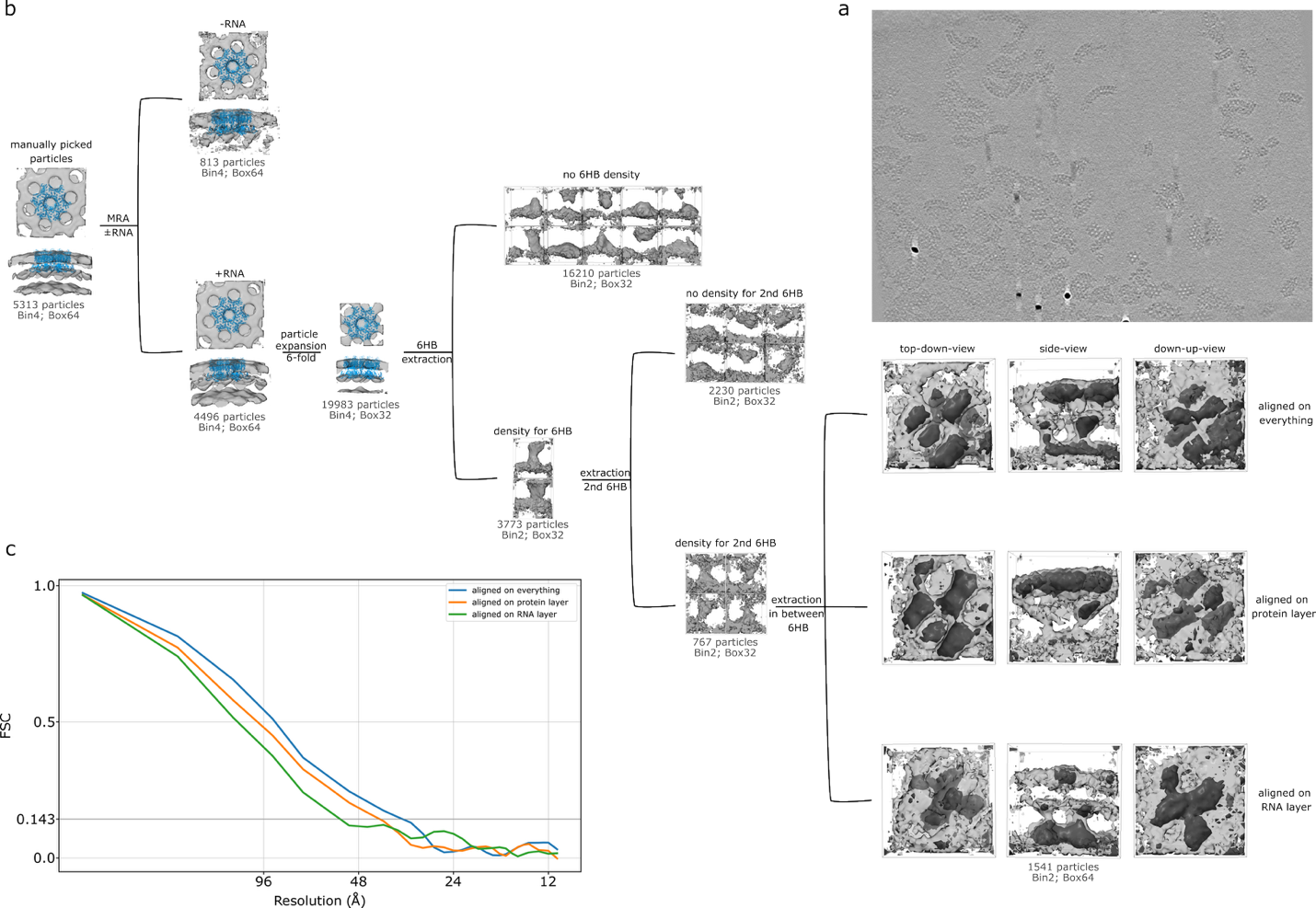

**Figure S8:** **Workflow and results of subtomogram averaging. a:** Representative tomogram used for structural analysis, showing higher molecular weight assemblies. **b:** Workflow of the single-particle tomography analysis. Manually picked particles were first aligned, then subjected to multi-reference classification to separate RNA-containing from RNA-lacking particles. To increase particle number, each particle was expanded to include the six adjacent CA hexamers. After realignment, particles were re-extracted in a smaller box size and screened for 6HBs connected to RNA density. This process was repeated for the adjacent 6HB, further enriching for RNA-associated assemblies. The final set of 657 particles was re-extracted, centered between the two 6HBs, and aligned either against the full density, the protein layer, or the RNA layer using appropriate masks. A merged model of PDB 4usn and 5i4t was fitted into the protein density (blue). Final structures are overlayed at two contour levels (gray and black). **c:** Fourier shell correlations (FSC) curves of the three final maps. Gold-standard resolution estimates (0.143 cutoff) were: overall alignment, 32.7 Å; protein alignment, 40 Å; RNA alignment, 51.5 Å (blue, orange, and green curves, respectively).

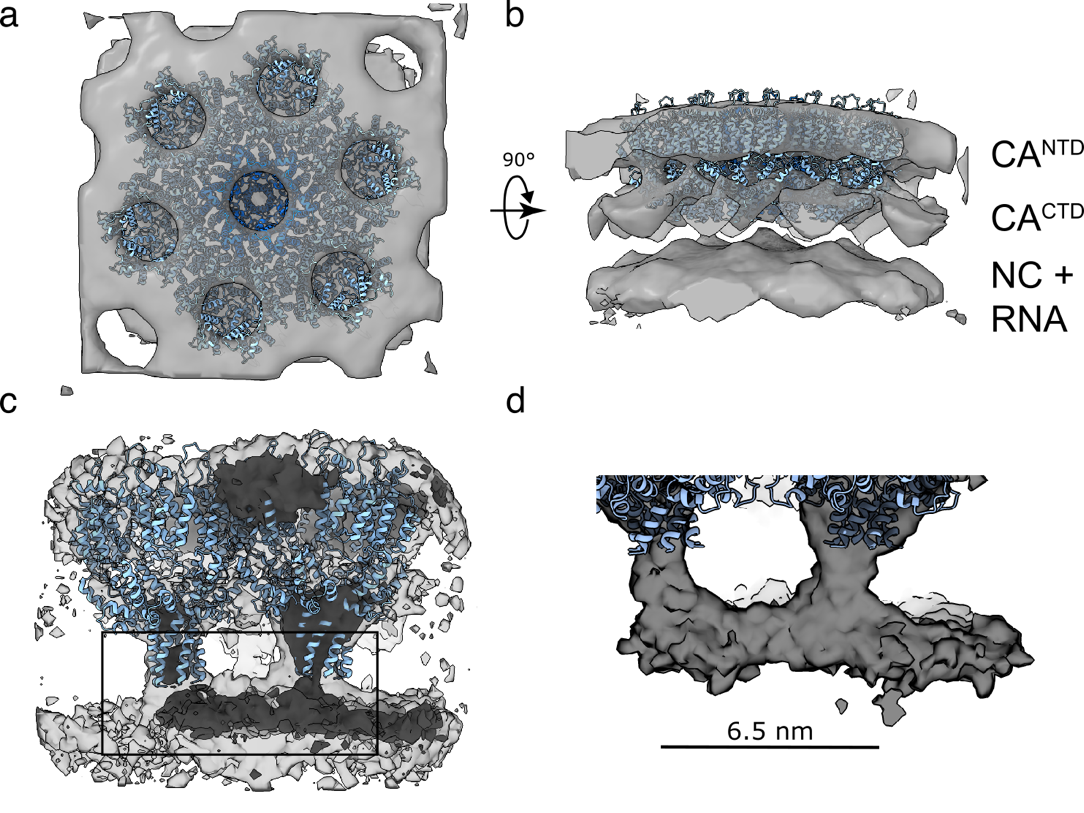

**Figure S9: Tubular RNA density underneath the protein layer of Gag assemblies formed with Ψ^CES^_2_ in single particle cryoET. a:** Initial reconstruction obtained by sub-tomogram averaging from manually picked particles aligned to the full assembly. The map reveals the hexameric capsid lattice. **b:** The same map rotated shows the two protein layers from CA^NTD^ and CA^CTD^, and an additional underlying density corresponding to NC-SP2-p6 and the Ψ^CES^_2_. **c:** Final reconstruction after particle selection for strong RNA-layer signal. A tubular density is observed bridging two adjacent 6HBs. The same map is displayed at two contour levels (high, light gray; low, dark gray). **d:** Zoomed view of the tubular RNA density of a length of 6.5 nm connecting the two central six-helix bundles. PDB entries 4usn and 5i4t were merged and fitted into the protein density (blue).

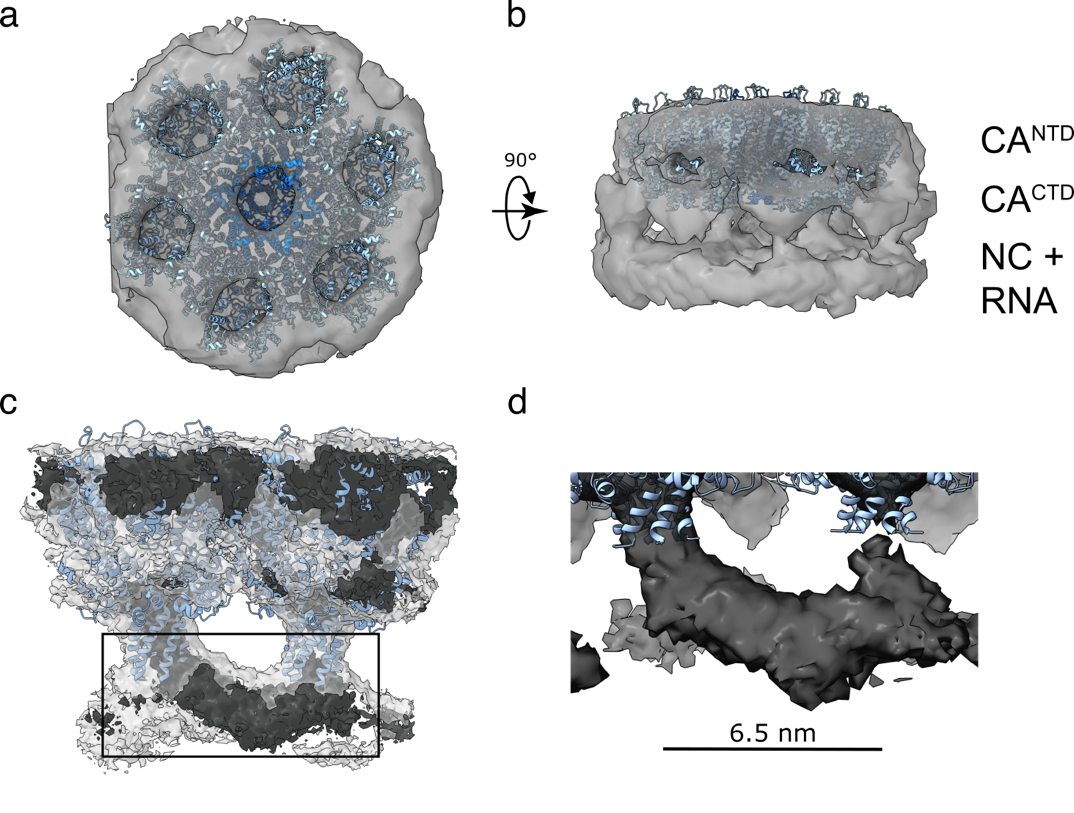

**Figure S10: Tubular RNA density underneath the protein layer of Gag assemblies formed with DIS_2_ in single particle cryoET. a:** Initial reconstruction obtained by sub-tomogram averaging from manually picked particles aligned to the full assembly. The map reveals the hexameric capsid lattice. **b:** The same map rotated shows the two protein layers from CA^NTD^ and CA^CTD^, and an additional underlying density corresponding to NC-SP2-p6 and the DIS_2_. **c:** Final reconstruction after particle selection for strong RNA-layer signal. A tubular density is observed bridging two adjacent 6HBs. The same map is displayed at two contour levels (high, light gray; low, dark gray). **d:** Zoomed view of the tubular RNA density of a length of 6.5 nm connecting the two central six-helix bundles. PDB entries 4usn and 5i4t were merged and fitted into the protein density (blue).

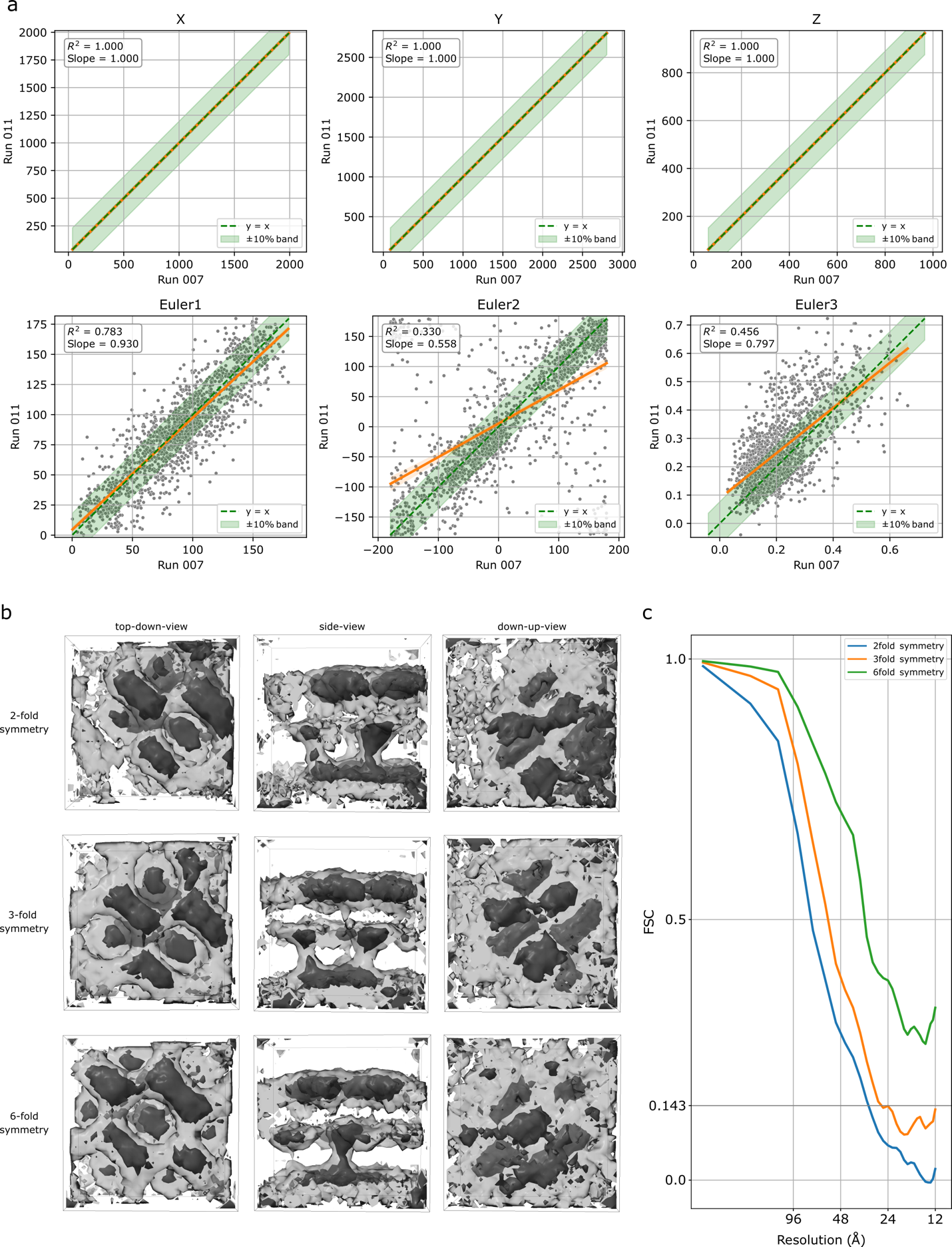

**Figure S11:** **Comparison of coordinate and angular alignments, and effect of symmetry on maps. a:** Scatterplots showing x, y, and z coordinates of particles aligned on the protein layer (x-axis) versus the RNA layer (y-axis) (top three plots). The absence of systematic deviations indicates no translational bias between layers. By contrast, plots of the three Euler angles (bottom three plots) reveal differences, suggesting rotational heterogeneity between the two layers. A 10% interval range is highlighted in light green; the x=y line is shown as a dotted green line, and the regression line as orange. **b:** Maps calculated with the final subset of 657 particles using 2-fold, 3-fold, or 6-fold symmetry. The overall appearance of the maps is similar compared to C1; however, RNA density (tubular features) weakens progressively with higher symmetry, becoming nearly absent under C6, while the hexameric protein lattice remains robust. **c:** Gold-standard resolution estimates (0.143 cutoff) were: 30 Å (2-fold symmetry; blue), 25.7 Å (3-fold symmetry; orange). For C6 (green), the FSC curve did not cross the 0.143 threshold.

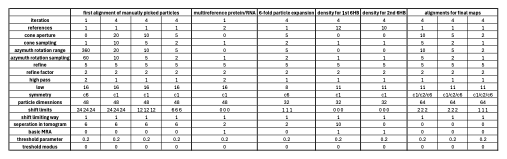

**Table S1: Specific refinement parameters used for each round of the single particle tomography data analysis.**
